# HUNGER DRIVES SWITCHING AND SEARCHING RESPONSE IN A SOCIAL PREDATOR

**DOI:** 10.1101/2023.11.27.568908

**Authors:** CM Prokopenko, S Zabihi-Seissan, DLJ Dupont, KA Kingdon, JW Turner, E Vander Wal

## Abstract

Hunger is a frequent state for many predators and increasing hunger is likely to motivate costly behaviour to acquire necessary resources. Generalist predators must balance the costs and gains of hunting different prey, including increasing encounter rates and improving success rates by seeking areas with greater prey catchability. Large carnivores face threats when they interact with humans or conspecifics. We use integrated step selection analysis to describe spatiotemporal factors that influence wolf (*Canis lupus*) hunting behavior in Riding Mountain National Park, a natural area that wolves share with moose (*Alces alces*) and elk (*Cervus canadensis*). If hunger generates more risky behavior by wolves, as time-from-kill increases we expect wolves will: (1) search for and kill a prey that poses higher risk of injury, (2) use the periphery of their range, (3) use areas closer to the park boundary. Hunger alters wolf space use and drives a fine scale change in prey tracking. Movement patterns of hungry wolves are indicative of search behavior, i.e., shorter steps and more turning. Contrary to our predictions, hungry wolves moved further into the park. As wolves become hungrier, they switch their response from a weak selection to avoidance of elk. In contrast, the response to the primary and emergent prey, moose varied between individuals with some pack level similarities. Therefore, the state-based response to a pervasive risk and a historical resource was conserved in a population residing in a prey rich ‘island’ interfacing with human disturbance.

## Introduction

Nearly half a century ago, Charnov’s (1976) germinal experiment depriving mantises of food changed the way we think about foraging theory; state-dependence, such as hunger, is now foundational in movement ecology (Nathan et al., 2008). Just as movement cannot often be disconnected from state, it also cannot be separated by the motivation to occupy a space. Habitat selection is the manifestation of evolved behaviors driven by anticipated energetic gains and cost of using a particular space. The emergent responses to the pressure to obtain resources and manage risk are offset or magnified by individual qualities, for example nutritional state (DeWitt et al., 2017).With large carnivores often experiencing low hunting success, hunger is likely a baseline state, and increasing hunger motivates risk-taking to acquire resources. For instance, hungry cougars (*Puma concolor*) were more likely to use human developed areas, which are normally avoided due to risk (Blecha et al., 2018). By incorporating state dependence into movement integrated habitat selection, we can therefore test how cooperative and cursorial hunters mediate spatial trade-offs.

As animals become hungrier, foraging behaviour tips in favour of risk-taking. Hungry barn owls (*Tyto alba*) perform a risky hunting behaviour by increasing their attacks (Embar et al., 2014). Risks are posed by the resources themselves (Mukherjee & Heithaus, 2013) and a more desperate predator may pursue more dangerous resources. Common starling (*Sturnus vulgaris*) consumption of toxic prey increased with decreasing energetic condition (Barnett et al., 2007). Some risky areas may become more worthwhile to hungry consumers if those areas co-occur with resources. Hungry spiders (*Pardosa milvina*) foraged when predator cues were present, while sated counterparts avoided the areas both high in risk and high in resources (Walker & Rypstra, 2003). The individual variation in risk-sensitive foraging has population implications (Sinclair & Arcese, 1995). Therefore, the ability of internal state to alter the balance between resource and risk is fundamental to descriptions of wildlife space-use.

Habitat selection and movement modelling are tools to describe how animals respond to external variation and can test expectations set by optimal foraging theory. Optimal foraging theory addresses how the energetic value of an area to an animal depends on the interplay of resource and risk (MacArthur & Pianka, 1966). The assumption that animals will spend more time in areas that confer higher fitness is fundamental to both habitat selection analysis and optimal foraging theory. Mortality risk reduces the value of an area and subsequently an animal’s time allocation in that area (Brown, 1999). Movement enables optimal foraging by allowing animals to mediate trade-offs in space and time. For example, elk spend more time in high-risk foraging areas when wolf activity is lower (Kohl et al., 2018). Since its conceptualization, optimal foraging theory has addressed the effect of hunger on space-use behaviour (Charnov, 1976). Habitat selection analyses can test optimal foraging theory and there is a recent proliferation of approaches to quantify energy landscapes (Berti et al., 2022; Klappstein et al., 2022). Here we explore the internal state-dependent habitat selection and movement behaviour of an apex predator.

Hunger is an internal state that changes over fine temporal scales. Integrated step selection analysis can incorporate fine scale temporal change to test how hunger influences wolf response to spatial trade-offs in Riding Mountain National Park. If increased hunger generates more risky behaviour by wolves to expand their hunting opportunities, as time-from-kill increases we expect wolves will: (1) search for and kill prey that pose higher risk of injury, (2) use areas on the edges or outside of pack ranges where the conflict with other packs is more likely and (3) be closer to high-traffic human use areas inside the park, or venture outside the park boundary, which increases the potential for human interactions.

## Methods

### Study area

Riding Mountain National Park (RMNP; 50°051′50″, N 100°02′10″W) is a 3,000 km^2^ conserved area in southwestern Manitoba. The conserved habitat (a confluence of prairie grassland, aspen parkland, and boreal forests) creates a distinct edge with surrounding agricultural land. During the study period (2016-2017), the wolf population was estimated from aerial and snow track surveys at ∼70 individuals and 13 packs. Mortality causes (68% of the sample population) observed during our study were anthropogenic (trapping, poisoning, gunshot; constituting 20% of mortalities of study animals), conspecific (12%), and disease (Canine Distemper Virus 36%; Turner et al., 2023). Prey defense is a source of mortality not represented in the sample population, but one unmarked individual was discovered at a kill site, succumbing to injuries due to blunt force. Aerial surveys to estimate ungulate population sizes were conducted by Parks Canada annually in the winter (Figure S1). Prey available to wolves include moose (*N* = 2,300 animals), elk (*N* = 1,100), and white-tailed deer (*Odocoileus virginianus;* N = 750). There was no evidence that livestock residing outside the park were actively hunted by our collared wolves, though some wolves occasionally and opportunistically scavenged bait stations and dump sites outside the park.

### GPS Location data

We collared 35% of wolves in the RMNP population and 60% of wolf packs; at least one wolf was collared in all packs in the ‘core area’ on the west side of RMNP (Figure S1). Wolves in winter 2016 (*n* = 13) and 2017 (*n* = 14) were captured by Bighorn Helicopters following Memorial University AUP 16-02-EV and fit with GPS telemetry collars (Advanced Telemetry Systems G2110E, MN USA; Followit Tellus Medium, Followit Sweden AB, Lindesberg, Sweden; Lotek Iridium TrackM 2D, Lotek Wireless Inc, Newmarket, ON, Canada; Sirtrack Pinnacle G5C, Sirtrack Limited, Hawkes Bay, New Zealand; Telonics TGW-4577-4, Telonics Inc., AZ USA). Location data was rarified to a two-hour interval to sample individuals at an equal intensity.

### Cluster Investigation

Continuous and extensive fieldwork investigations determined the timing and location of wolf behaviours, including wolf killed prey. Important areas of wolf activity were indicated by an increased density of GPS locations, i.e., ‘clusters’. Clusters were identified in Python from an version of the code presented in Knopff et al. (2009) and originally created by Warren (2008)adapted for wolves (DeCesare, 2012; Irvine et al., 2022; Webb et al., 2008). In our study, the inclusion rules were set to a radius of 300m and a time of 96 hours, meaning that if a new location was within those limits from any of the locations currently in the cluster it was added.

Once clusters were investigated, unique areas were then termed ‘sites’ based on the associated behaviours identified (detailed methods and results of cluster investigation are covered in Prokopenko, 2022). Often multiple clusters occurred at each unique site and multiple wolves visited the same site. A total of 1260 clusters were investigated, which translated to 598 unique sites. Aggregate clusters of wolves were defined from spatial and temporal to determine unique kills. The primary behaviour was ‘kill’ at 181 unique sites (433 clusters designated as kills), probable kills at 24 sites, and scavenge at 46 sites. A total of 296 scat samples were collected and subsequently analyzed. Both kill site and scat data indicated moose contributed over half of the biomass in wolf diets (65% kill sites, 55% scat). Elk contributed less than half of the biomass (21% kill sites, 34% scat).

### Integrated step selection analysis

We conducted the movement and selection analysis for individual wolves during the winter period, January to March, resulting in an average of 190 locations per wolf-winter (Table S1). Lone wolves demonstrating extraterritorial movements outside the park or wolves without kills identified during the winter were omitted from the analysis to promote model convergence leaving 21 wolves from seven packs in this analysis.

Integrated step selection analysis (iSSA) estimates movement and selection behaviour together to reduce bias and produce a holistic depiction of factors that motivate and modify animal space-use (Avgar et al. 2016). We used iSSA to incorporate fine-scale temporal-dependence into our models.

Used step lengths (distance between two consecutive GPS locations) were described by a gamma distribution (mean tentative shape = 0.419, scale = 218, Table S1). The directionality of steps was defined by turn angles (angular deviation from the step heading) fit with a Von Mises distribution (mean tentative kappa = 0.157, Table S1). Observed movement behaviour informed the availability domain. We randomly sampled step lengths and turn angles from these distributions to create ten available steps for each used step. Within each stratum of 11 steps, the start points are shared but the end points are distinct. Movement and selection covariates at the start point are those thought to influence the subsequent space-use decision, while those included at the end point estimate the resulting selection pattern of an animal. When step length is ln-transformed (natural log) and included in a step selection model, the resulting covariate is a modifier of the shape parameter of the original gamma distribution. The cosine of the turn angle transforms this circular measure to a linear correlation with previous step heading (-1 is backward movement, 1 is forward). In a step selection analysis, the coefficient relates to the concentration parameter of the Von Mises distribution.

We generated five spatial layers of covariates to describe risk and resources experienced by RMNP wolves. We calculated the distance from the boundary of RMNP (shapefile from Parks Canada), areas outside of the boundary were given a value of ‘0’. Only two main roads exist in the park, compared to a dense network outside. Thus, distance to road was not included in the model as it is highly correlated with the park boundary and risky areas. Risk from conspecifics was estimated as a continuous variable using distance to range centre. To estimate the centre of the wolf pack territories, we used 90% minimum convex polygons (MCPs) around all locations from individuals in each pack.

Prey catchability was calculated using a habitat selection framework where ‘used’ points were kill site locations determined from kill site investigation and available points were drawn uniformly within each pack range. Covariates included in these models were landcover, distance to water, distance to roads, distance to maintained and unmaintained trails, distance to hard edge, and terrain ruggedness. Hard edge was calculated using the transition zone between open areas and closed canopy forest. Prey catchability layers were created for the three ungulate prey species in RMNP: elk, moose, and white-tailed deer. Models for elk and moose were conducted for separate years and matched to the wolf-year. Deer kills occurred mostly in 2017, thus, these year differences could not be accounted for. White-tailed deer kills increased in mixed wood (Table S2). Generally, moose kills increased in mixed wood and areas closer to maintained trails and elk kills increased closer to unmaintained trails and edge (Zabihi-Seissan et al., 2022).

We calculated the time from leaving the last known kill for each location as an estimate of hunger. Time from kill at the start point of the cluster was included as an interaction with the movement and selection covariates. Locations that contributed to the cluster were removed from the data used in the iSSA so that inferences reflect space-use behaviour when predators are pursuing (travelling steps) or conserving (resting sites) energy. A similar designation of hunger measured was used on cougars, specifically time since feeding activity binned into 11 intervals from 0-1 days to over 10 days (Blecha et al. 2018). In our analysis we used a continuous linear measure of hunger until 2 weeks, at which point we censored data as our confidence in no kill events occurring over these long intervals decreased (20% of the data was censored using this rule).

Step length and turn angle estimation, random step creation, and covariate extraction was completed using the *amt* package (Signer et al., 2019) in R version 4.1.0. We fit a mixed effects iSSA with individual random slopes for covariate interactions with time from kill using the *glmmTMB* package in R to fit a conditional Poisson model (Muff et al., 2020).

### Calculating effect sizes

Relative selection strength (RSS, Avgar et al., 2017) provided an illustration of the effect size of selection coefficients. Specifically, we demonstrated the change in selection for prey catchability, pack range centre, and the park boundary with time from kill. We used the *predict* function to compare the selection of one location of average habitat over another location that has high values of the covariate of interest. We held the difference in the location habitat covariates constant while varying time from kill (0 to 7 days from kill). We used individual-specific random effects to present uncertainty and individual variation around population responses.

The change in speed as time from kill increased illustrated the effect of hunger on movement. We calculated speed (meters per hour) from the mean step length of the gamma distribution (meters), divided by the fix interval (2 hours). The mean step length was the shape parameter multiplied by the scale. We modified the shape parameters estimated for the random steps by the coefficients that included ‘ln Step Length’.

## Results

Wolf spatial behaviour changed with hunger (days from kill). Wolves selected for the catchability of their ungulate prey, but these responses were sensitive to their hunger. Wolves had a neutral response to moose catchability (0.034 [-0.521, 0.589]), which was not affected by hunger (0.044 [-0.161, 0.073], Figure 1a). Wolves weakly selected elk catchability (0.138 [- 0.288, 0.564]), this response switched to avoidance with increasing time from kill (-0.164 [- 0.292, -0.037], Figure 1b). Deer catchability was selected (0.957 [0.501, 1.41]), with a variable response as time from kill increased (-0.034 [-0.091, 0.160], Figure 1c). The variation in these responses were greatest for moose, followed by white-tailed deer, and the response was consistent for elk (Figure 1a-c).

**Figure 1.**
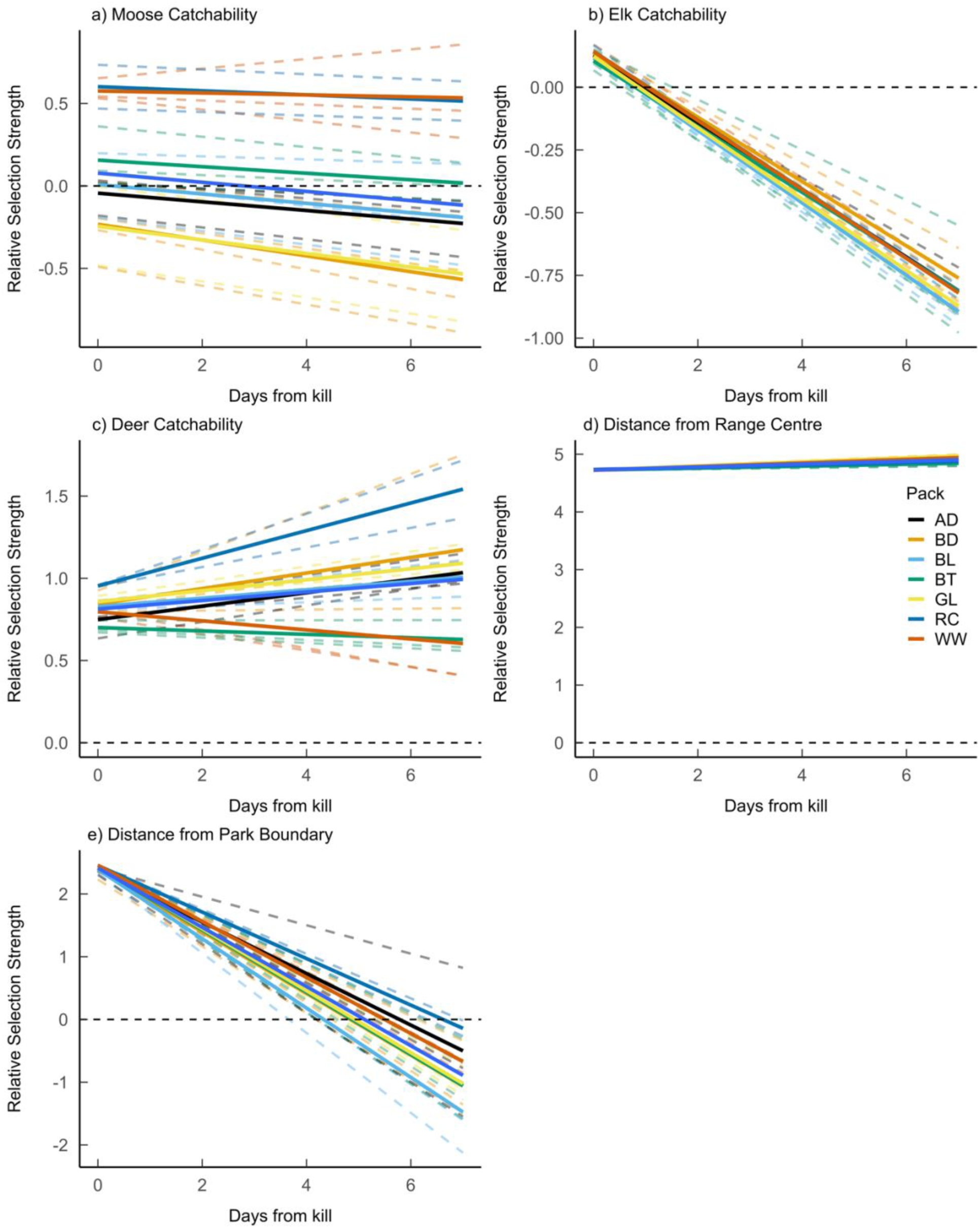
Natural log-transformed Relative Selection Strength (ln-RSS) for location x_1_ over another location x_2_ as time from kill increases (0 to 7 days). The ln-RSS for individual wolves is displayed as dashed lines coloured by pack. The dashed line at zero indicates no difference in selection between the two locations. The two locations are identical (including time from kill) except for differences in the value of a single habitat variable a) max (x_1_=1) versus mean estimate (x_2_=0.253) in moose catchability, b) max (x_1_=1) versus mean estimate (x_2_=0.137) in elk catchability, c) max (x_1_=1) versus mean estimate (x_2_=0.142) in deer catchability, d) minimum (x_1=_ 0 km) versus mean estimate (x_2_=8.61 km) from the centre of the corresponding pack range, e) minimum (x_1 =_ 0 km) versus mean estimate (x_2_=8.42 km) from the boundary of Riding Mountain National Park. Positive values indicate selection for x_1_.

The spatial response to areas of risk also varied with hunger. Wolves selected to be closer to their range centre (-0.549 [-0.695, -0.403]), which did not change as days from kill increases (0.004 [-0.033, 0.025]); Figure 1d). Wolves selected to be closer to the park boundary (-0.298 [- 0.396, -0.201]; then selected areas farther from the boundary as they became hungrier (0.056 [0.027, 0.085], Figure 1e). The response to pack range and park boundary had little variation between individuals population (Figure 1).

There is some evidence that wolf movement response was influenced by hunger. Wolves moved slower and less directionally than the estimated used to generate movement distributions for the available steps. The shape parameter of the gamma distribution of step length was lower than the available distribution as indicated by a negative ln-transformed step length coefficient (- 0.0487 [-0.122, 0.025]) and decreased with hunger (-0.006 [-0.015, -0.003], Figure 2a). A negative coefficient is interpreted as wolves taking more short steps. The coefficient for cosine of the turn angle was also weakly negative (-0.070 [-0.205, 0.346]) indicating wolves were less directional than the tentative kappa parameters (concentration of the Von Mises distribution of turn angle) estimated for the available steps. However, the estimates varied across the population and only was weakly influenced with increasing days from kill (-0.022 [-0.481, 0.005]; Figure 2b). The differences in movement varied by individual with similarities between pack (Figure 2).

**Figure 2.**
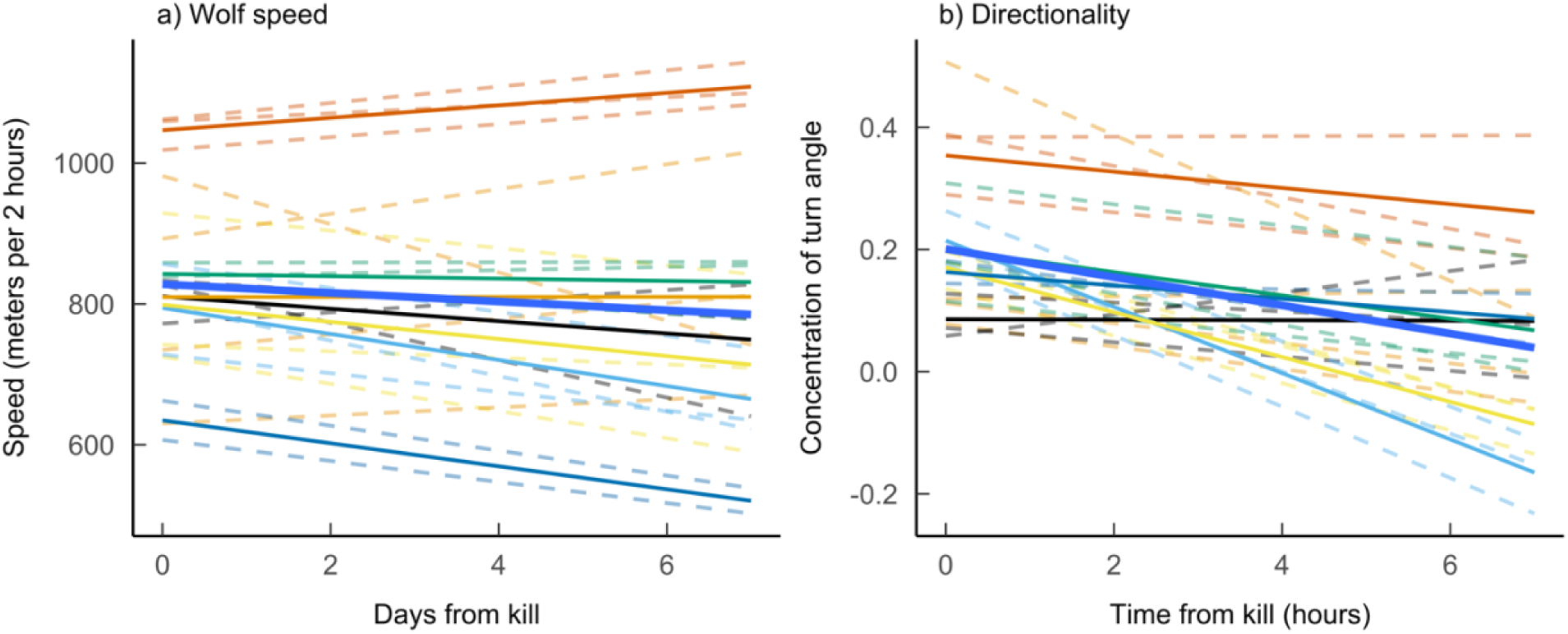
Movement response (speed in meter per 2 hours and concentration of turn angle) of wolves as a function of time from the final point in the previous kill cluster (0 to 7 days). Speed is calculated from the tentative shape and scale parameters of the gamma distribution of step length and modified by ln-transformed step length coefficients from the model output. Concentration of the turn angle is calculated by modifying kappa, the parameter from the Von Mises distribution. Cosine of the turn angle varies from -1 (indicating complete turns backwards) to 1 (indication continuing straight forward).

## Discussion

Hungry predators adjust their space use behaviour. Individual responses of wolves to longstanding specific risks and resources were aligned but then varied for many emergent or dynamic factors. Risk can come in several forms including the risk of injury while targeting large prey, lethal encounters with conspecifics, and human conflict. Predators strongly responded to the most pervasive risk: interactions with humans outside of the park. Wolves increased avoidance of areas closer to the park boundary with increasing hunger. Wolves switched their spatial response from selecting their preferred and historical prey, elk, when satiated to avoiding as hunger increased. Therefore, hunger mediates the trade-off between energetic reward and mortality risks.

Early movement pattern studies offer guidance to understand a predator’s response to heterogeneous resource distribution (Turchin, 1996). Specifically, predators are expected to spend more time in areas of high resource density (Fauchald & Tveraa, 2006). Area-restricted search behaviour corresponds with decreased movement rates and more torturous paths when foraging (Kareiva & Odell, 1987). There is some evidence that wolves in the study area are increasing their search effort when hungry by reducing the number of long steps and straightforward steps. Foundational prey theory (*i.e*., law of mass action and functional response) and empirical evidence supports the expectation that increased speed increases encounter and kill rates (Vander Vennen et al., 2016). Both these patterns can be occurring, as step selection analysis and the law of mass action operate under different assumptions. We modelled the spatially biased movement behaviour of wolves at a 2-hour time scale, which does not meet the assumptions of instantaneous measures in a homogenous system from the law of mass action. Both approaches are valid and can be synthesized to understand the spatial and temporal mechanisms of wolf foraging responses with changes in state.

Hungry wolves moved farther inside the park. Human activity poses mortality risk to predators and are often avoided(A. F. Smith et al., 2022). Cougars increased their selection for housing density when hungry, indicating energy depletion drove them to areas of higher payoff that simultaneously posed higher risk (Blecha et al., 2018). In this population of wolves, as hunger increased, they selected for greater distance from the park boundary. Notably, it was extremely rare for wolves to leave the safety of the park (>75% of wolf locations are within the park, Figure S2). When wolves were outside of the park their increased speeds were not indicative of a hunting behaviour (Figure S3). In addition, wolves selected to be closer to their range centre, which did not considerably shift with hunger. Here, we provide evidence that hunger is not driving human-wildlife interactions through increased spatial overlap for wolves in RMNP. In conclusion, hunger has the potential tilt the risk-reward trade-off towards more risky areas of the landscape, *if* those areas have higher energetic reward.

Hungry wolves shifted their selection from prey catchability, following a trade-off between ease of capture with probability of encounter. Predators exhibit varied strategies to improve hunting success including: tracking areas of abundance or habitat to encounter prey; selecting for areas that facilitate movement; and focusing efforts on prey that are more vulnerable or areas that increase prey capture (Balme et al., 2007; Kittle et al., 2017). This population of wolves select for prey spatial catchability at the within-territory scale (Zabihi-Seissan et al., 2022). In their territories, wolves had a strong positive selection response to moose habitat and moose catchability interaction (Zabihi-Seissan et al., 2022). At the step-level, satiated wolves strongly selected for deer, weakly selected for elk catchability, and had a variable response to moose catchability. Hungry wolves demonstrated a switch in their spatial prey tracking as the selection for elk decreased. The response to deer and moose was variable between individuals and packs. Therefore, spatial behaviour of wolves aligns with the observed population-level changes in wolf kill rates and prey abundance in this study area.

The variation in fine-scale spatial behaviour corresponds to the environmental pressures acting on the wolf population in RMNP. The kill site investigations that occurred during this period revealed a population-level switch from elk to moose. Our results indicate a similar spatial switch from elk; there was a strong population-level change with hunger was for elk. Further, deer abundance is increasing but distribution is concentrated to southern edges and closer to human disturbance (Zabihi-Seissan, 2019). Multiple predators have been documented shifting their space use in response to emergent and ephemeral prey (Svoboda et al., 2019). The variation observed in wolf space use response to changes in prey could indicate adapting behaviour to new environmental conditions, specifically prey abundance and distribution. If foraging behaviour is a labile trait, it is under the hungry condition that there is more variance, which generates opportunity for selection.

Cohesion of social groups emerges from a balance of risks and rewards (Krause et al., 2002; Silk, 2007). In our results, pack similarities emerge from individual wolf responses to the environment. Social carnivores are more cohesive when hunting larger and more dangerous prey (MacNulty et al., 2014; Smith et al., 2008). The study population of wolves in RMNP are more cohesive in areas used by larger prey and at kill sites of large prey (Zabihi-Seissan, 2019). These social constraints are likely contributing to the notable variation in patterns by pack for selection for moose catchability, the more abundant and risky prey, but not elk. Further, groupmates aid in defense of resource loss from scavengers or mortality risk from other large predators (Caraco & Wolf, 1975; Vucetich et al., 2004). Packs share similar genetics and environmental conditions, which both influence space-use. Owls in the same nest, regardless of genetic relatedness were more similar (Bombieri et al., 2018). The processes generating the phenotypic patterns in selection and movement within and between wolf packs should be investigated further.

Habitat selection manifests evolved behaviors driven by anticipated rewards and risks of using a particular location in space. By incorporating state-dependence into movement integrated habitat selection we illustrate how individual cooperative hunters have evolved to acclimate to increased risk as they become hungrier. This work adds to the building evidence that energetic trade-offs shape spatial responses (Berti et al., 2022; Klappstein et al., 2022) with added detail that hunger mediates those choices (Berger-Tal et al., 2009; Blecha et al., 2018; Embar et al., 2014). Overall, energy requirements and state can influence population distributions and survival as desperation makes risks more appealing.

**Table 1.**
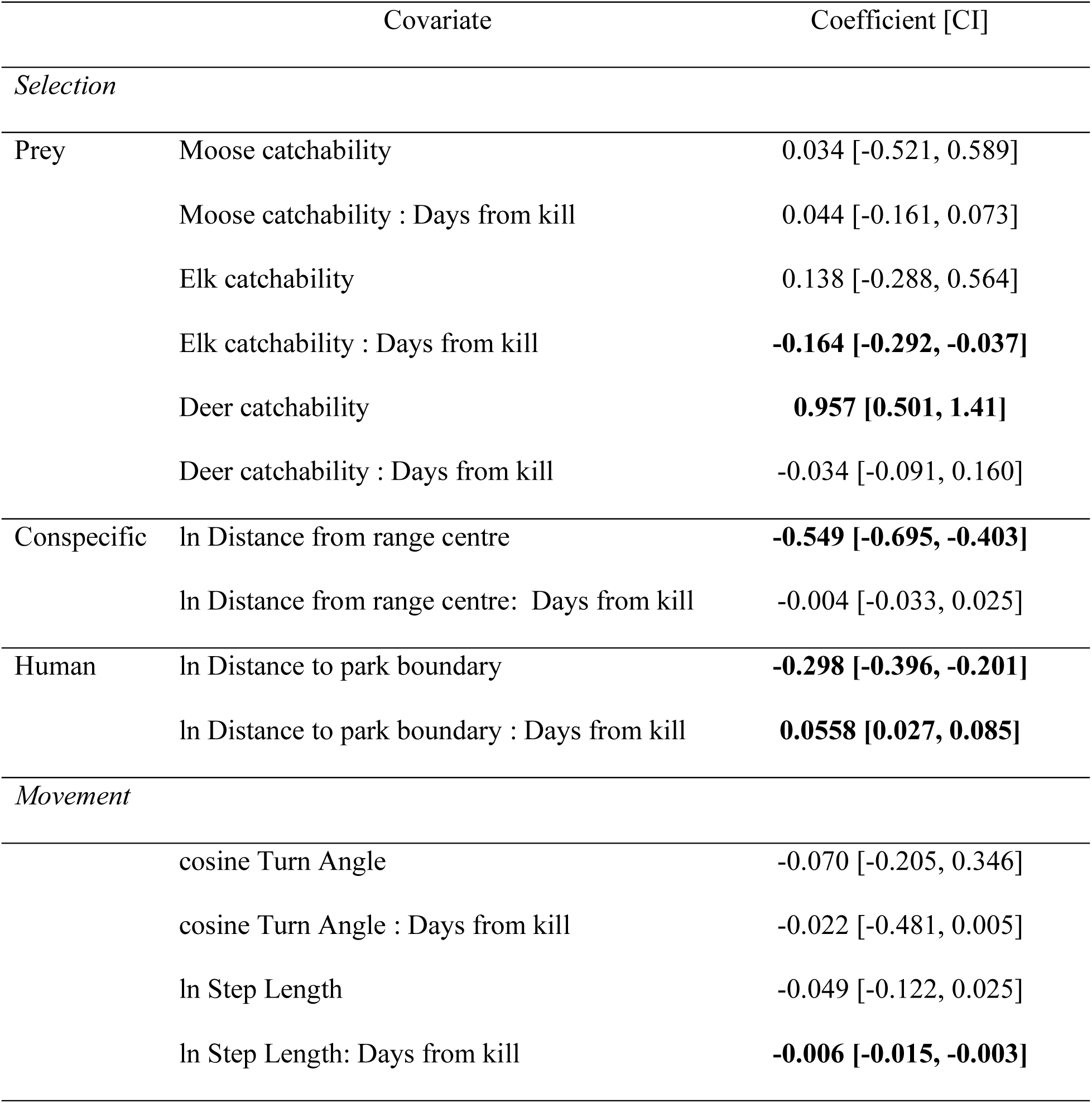
Population-level coefficients from the integrated step selection model fit with individual-specific differences (Muff et al., 2020) testing the change in wolf selection of resource or risk and movement behaviour with increasing hunger (days from kill). Bold coefficients indicate confidence intervals that do not overlap zero.

