## Supplementary figures and tables for "HUNGER DRIVES SWITCHING AND SEARCHING RESPONSE IN A SOCIAL PREDATOR": Prokopenko-hungrywolf- SupportingInformation.docx

### Supporting Information

#### Input Data and iSSA parameters

**Table S1**. Summary table of collared wolf ID (bold indicates they were included in the analysis, n = 21), pack ID, used locations (in a strata with 10 available points), tentative gamma shape, scale, kappa shape. Individual coefficients from the model provided as supporting data.

| Wolf ID | Pack ID | Collaring Date | Used Locations  (Jan– Mar) | Shape | Scale | Kappa |
| --- | --- | --- | --- | --- | --- | --- |
| **01** | Gunn Lake | 18/Jan/16 | 118 | 0.36132237 | 2197.12192 | 0.05475558 |
| **02** | Whitewater | 20/Jan/16 | 349 | 0.3962194 | 2651.77314 | 0.28421483 |
| **03** | Baldy Lake | 21/Jan/16 | 314 | 0.33994584 | 2483.7291 | 0.07652301 |
| **04** | Baldy Lake | 19/Jan/16 | 297 | 0.29521074 | 2499.6679 | 0.12905542 |
| **05** | Gunn Lake | 20/Jan/16 | 62 | 0.55817801 | 1836.46041 | 0.12612302 |
| **06** | Whitewater | 18/Jan/16 | 307 | 0.61605537 | 1745.5957 | 0.29198696 |
| **07** | Baldy Lake | 14/Mar/16 | 47 | 0.44280656 | 2586.38024 | 0.45916766 |
| 08 | Lone Wolf | 26/Jan/16 | 0 |  |  |  |
| 09 | Lone Wolf | 27/Jan/16 | 0 |  |  |  |
| **10** | Baldy Lake | 27/Jan/16 | 297 | 0.47131447 | 1970.92513 | 0.12739392 |
| **11** | Gunn Lake | 20/Jan/16 | 142 | 0.49732863 | 1681.45041 | 0.16125508 |
| **12** | Whitewater | 20/Jan/16 | 299 | 0.54819785 | 1981.95376 | 0.23867235 |
| 13 | Gunn Lake | 14/Mar/16 | 0 |  |  |  |
| **14** | Lake Audy | 14/Feb/17 | 529 | 0.35109155 | 2440.52882 | 0.07622143 |
| **15** | Block | 13/Feb/17 | 240 | 0.40626602 | 2168.37597 | 0.14995776 |
| 16 | Spruce Lake | 13/Feb/17 | 0 |  |  |  |
| **17** | Ranch Creek | 13/Feb/17 | 216 | 0.30702876 | 2271.3429 | 0.15828021 |
| **18** | Ranch Creek | 13/Feb/17 | 203 | 0.31408058 | 2348.83172 | 0.09051765 |
| **19** | Birdtail Valley | 14/Feb/17 | 298 | 0.36740815 | 2642.2791 | 0.15068368 |
| **20** | Lake Audy | 14/Feb/17 | 286 | 0.40987091 | 2033.40613 | 0.0092104 |
| 21 | Spruce Lake | 13/Feb/17 | 0 |  |  |  |
| **22** | Birdtail Valley | 14/Feb/17 | 239 | 0.45398585 | 2052.52099 | 0.06714728 |
| 23 | Spruce Lake | 13/Feb/17 | 0 |  |  |  |
| **24** | Block | 14/Feb/17 | 180 | 0.51934873 | 1562.19645 | 0.15046035 |
| **25** | Block | 13/Feb/17 | 197 | 0.33823499 | 2994.93348 | 0.2313691 |
| **26** | Birdtail Valley | 14/Feb/17 | 238 | 0.3571711 | 2461.43977 | 0.22878853 |
| **27** | Lake Audy | 14/Feb/17 | 308 | 0.44632291 | 1973.39982 | 0.03559697 |

**Table S2.** Deer catchability model output from a logistic regression analysis (binomial, logit) where kills were designated ‘1’ and available ‘0’ were drawn across wolf territories. Outputs and detailed methods for the moose and elk catchability model are reported in Zabihi-Seissan et al. 2022. Supplementary Material Table S4. Significant values in bold

| **Term** | **Coefficient Estimate** | **Std. Error** | **P-value** |
| --- | --- | --- | --- |
| Intercept | -5.1689056 | 3.7284546 | 0.16564 |
| ConBog | 1.1409895 | 1.6384719 | 0.48619 |
| MarshGrass | 0.5299321 | 1.4752452 | 0.71943 |
| **Mixedwood** | **2.5498129** | **0.8698398** | **0.00337** |
| log_BTrail_Dist | -0.1949736 | 0.1537350 | 0.20471 |
| log_Road_Dist | 0.1500584 | 0.2891781 | 0.60382 |
| log_ Trail_Dist | -0.0355165 | 0.1857109 | 0.84833 |
| log_Water_Dist | 0.2035007 | 0.3027018 | 0.50140 |
| log_Edge_Dist | -0.1049687 | 0.1476021 | 0.47699 |
| log_Stream_Dist | -0.2620656 | 0.1446049 | 0.06994 |
| Ruggedness | -0.0004991 | 0.1066569 | 0.99627 |

**Table S3.** Model performance was evaluated using the performance package, check_collinearity were determined to have Low Correlation, check_overdispersion reported no dispersion (dispersion ratio = 0.872, Pearson's Chi-Squared = 49639.122, p-value = 1). VIFs for terms in the model are reported in the table.

| **Term** | **VIF** | **Increased SE** | **Tolerance** |
| --- | --- | --- | --- |
| log_sl | 1.13 | 1.06 | 0.88 |
| cos_ta | 1.01 | 1.01 | 0.99 |
| elk_kill_end | 2.15 | 1.47 | 0.47 |
| moose_kill_end | 1.39 | 1.18 | 0.72 |
| deer_kill_end | 2,02 | 1.42 | 0.50 |
| log_wolf_dist_end | 1.64 | 1.28 | 0.61 |
| log_park_dist_end | 1.59 | 1.26 | 0.63 |
| log_sl : tfkill_days | 1.12 | 1.06 | 0.89 |
| cos_ta : tfkill_days | 1.00 | 1.00 | 1.00 |
| tfkill_days : elk_kill_end | 2.14 | 1.46 | 0.47 |
| tfkill_days : moose_kill_end | 1.45 | 1.20 | 0.69 |
| tfkill_days : deer_kill_end | 2.10 | 1.45 | 0.48 |
| tfkill_days : log_wolf_dist_end | 1.84 | 1.36 | 0.54 |
| tfkill_days : log_park_dist_end | 1.65 | 1.29 | 0.60 |

#### Maps and Figures of wolves in RMNP

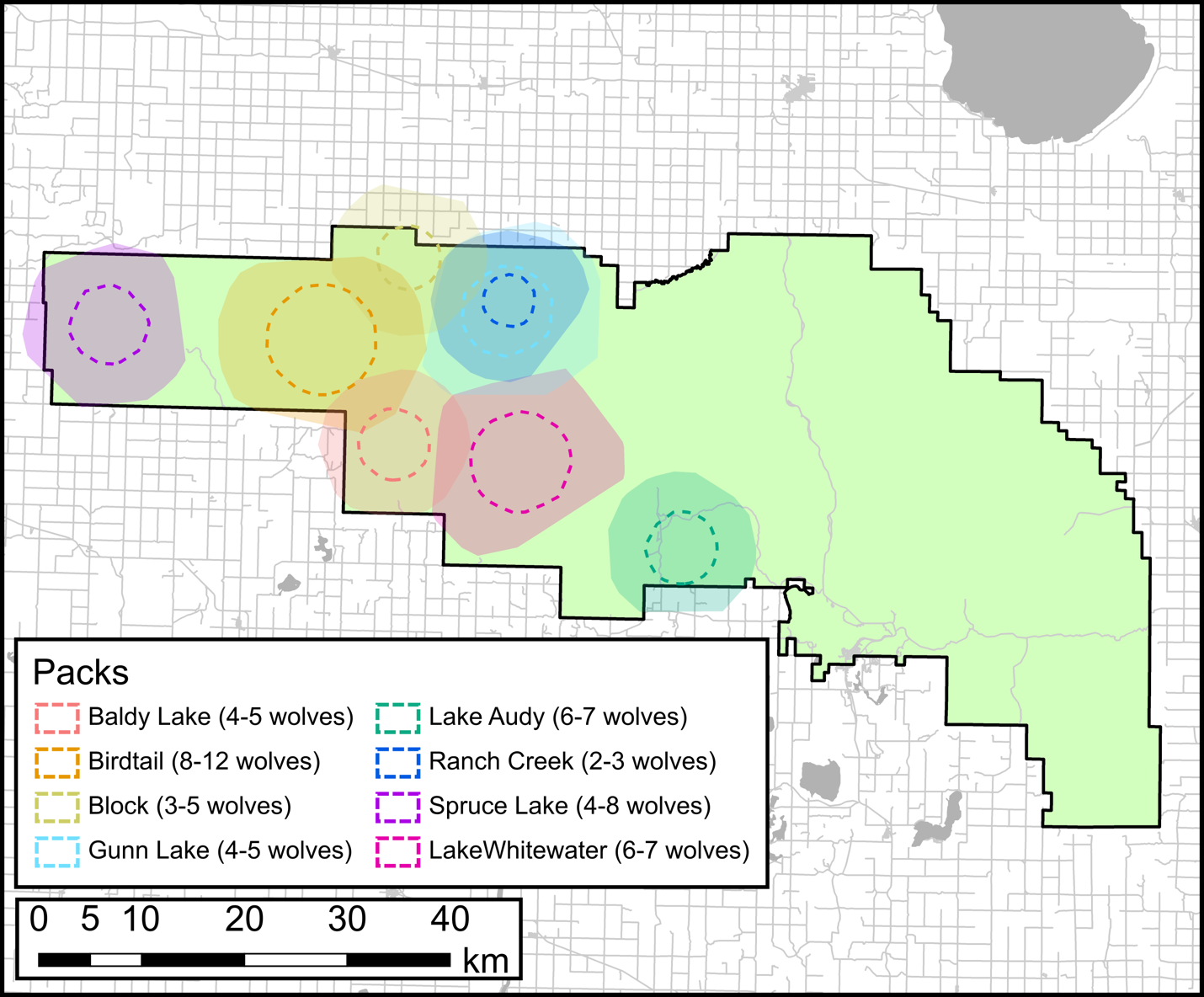

**Figure S1.** Park map with wolf ranges from collars on the west core area of Riding Mountain National Park. Pack home ranges and estimated number of wolves in each pack based on aerial

visual observations and trail camera photos. Shaded areas consist of tradition 95% minimum convex polygon which contain 95% of all wolf GPS points while the dotted lines consist of the core home range (50% minimum convex polygon). The W11 from Gunn Lake 2016 pack and lone wolf W09 established the Ranch Creek pack in 2017.

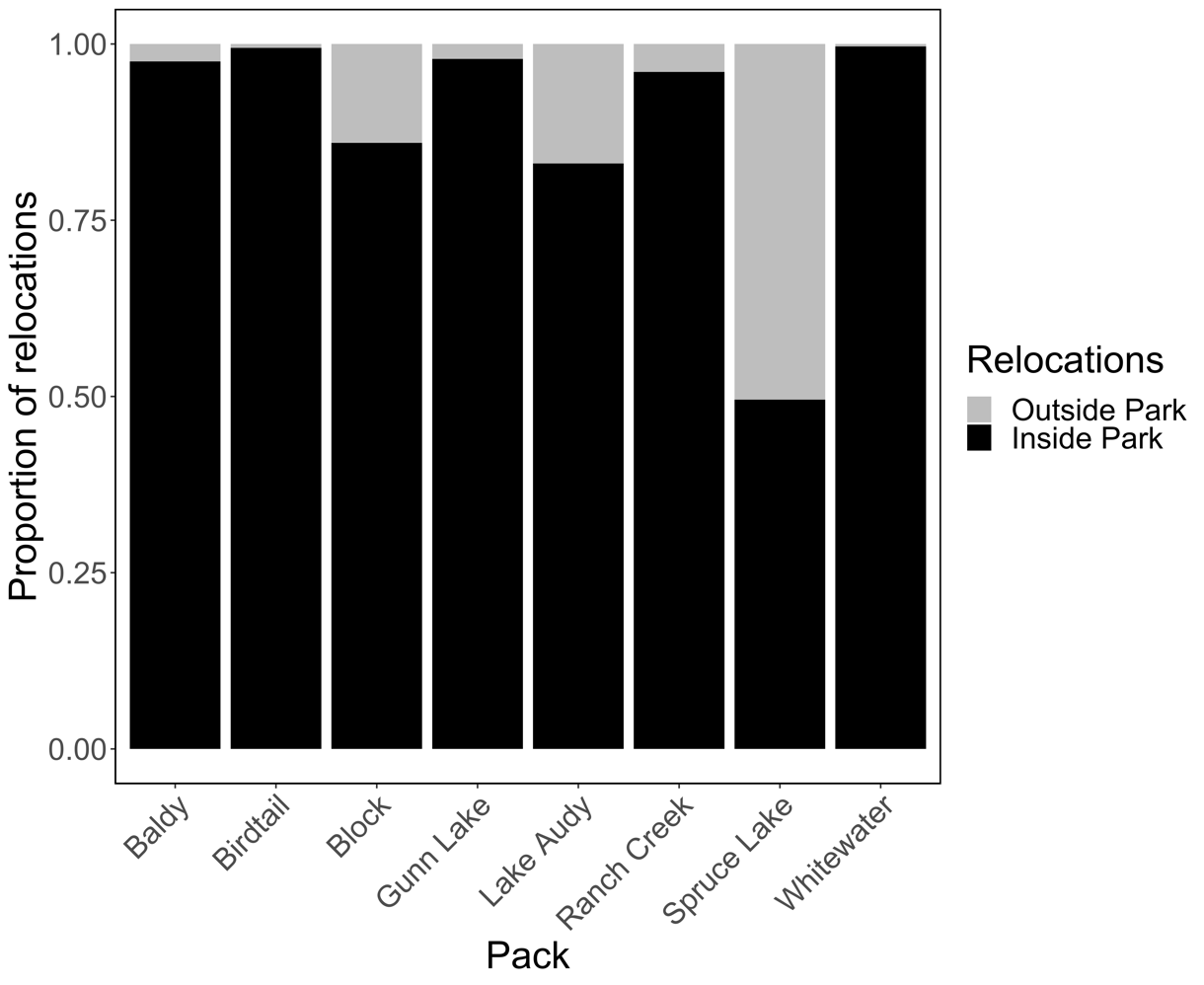

**Figure S2.** Proportion of locations inside and outside the for all collared wolves by pack 2016-2017.

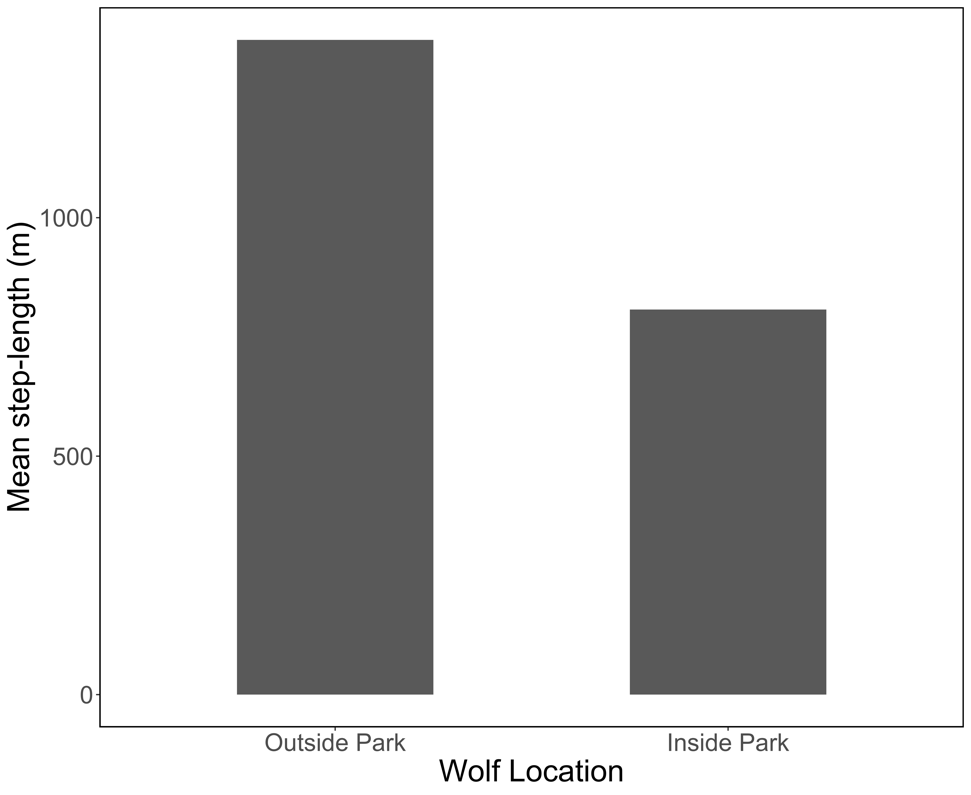

**Figure S3.** Step lengths of locations inside the park and outside the park for all collared wolves 2016-2017.
